# Critical nodes reveal peculiar features of human essential genes and protein interactome

**DOI:** 10.1101/831750

**Authors:** Alessandro Celestini, Marco Cianfriglia, Enrico Mastrostefano, Alessandro Palma, Filippo Castiglione, Paolo Tieri

## Abstract

Network-based ranking methods (e.g., centrality analysis) have found extensive use in systems biology and network medicine for the prediction of essential proteins, for the prioritization of drug targets candidates in the treatment of several pathologies and in biomarker discovery, and for human disease genes identification. We here studied the connectivity of the human protein-protein interaction network (*i.e.*, the *interactome*) to find the nodes whose removal has the heaviest impact on the network, *i.e.*, maximizes its fragmentation. Such nodes are known as Critical Nodes (CNs). Specifically, we implemented a Critical Node Heuristic (CNH) and compared its performance against other four heuristics based on well known centrality measures. To better understand the structure of the interactome, the CNs’ role played in the network, and the different heuristics’ capabilities to grasp biologically relevant nodes, we compared the sets of nodes identified as CNs by each heuristic with two experimentally validated sets of essential genes, *i.e.*, the genes whose removal impact on a given organism’s ability to survive. Our results show that classical centrality measures (*i.e.*, closeness centrality, degree) found more essential genes with respect to CNH on the current version of the human interactome, however the removal of such nodes does not have the greatest impact on interactome connectivity, while, interestingly, the genes identified by CNH show peculiar characteristics both from the topological and the biological point of view. Finally, even if a relevant fraction of essential genes is found via the classical centrality measures, the same measures seem to fail in identifying the whole set of essential genes, suggesting once again that some of them are not central in the network, that there may be biases in the current interaction data, and that different, combined graph theoretical and other techniques should be applied for their discovery.

## I. Introduction

Given a graph we say that it is connected if there is a path between each pair of vertices, and the graph *connectivity* is defined as the number of connected pairs. In this context the most important nodes of the network are those whose removal disconnect the graph *i.e.*, those who separate it into isolated subgraphs. The definition of connectivity is strongly related to network resilience, the ability of the network to resist to node removals. More generally, depending on the network topology, nodes with high connectivity can be related to different roles such as the spreading of news in social networks or the inhibition of virus diffusion.

More specifically, we addressed the problem of identifying the nodes whose removal result in a maximum fragmentation of the network. This problem has been studied by Borgatti in [7], formalized by Aruselvan *et al.* in [3] and in literature it is commonly referred to as the Critical Nodes Problem (CNP)^1^. The CNP defines as *critical nodes* (CNs) the set of nodes whose removal results in the maximal reduction of the number of connected node pairs. CNP is proven to be a NP-complete problem [3] and thus there are not algorithms known that can solve it in polynomial time. Practically, there are not computer programs known that can exactly solve it in a reasonable amount of time even for small graphs with hundreds of nodes. As a consequence, practical heuristics must be developed to solve CNP in real networks. As we suppose that CNs are *important* nodes in the network, a possible approach could be to rank nodes by a well known centrality measure (like the betweenness centrality), remove the node from the network and compute the remaining network connectivity. The choice of the measure depends on the phenomena to be analysed [7], [33].

The CNP can be of interest in network biology, thus in this work we focused our attention on the connectivity and topology of the human protein-protein interaction (PPI) network, or *interactome*. The understanding of the topology of the human interactome could have a tremendous impact on the comprehension of the molecular ground of human diseases. Indeed, the PPI framework is an integrative approach [13], [22], [25], [28], [31] that has already been extensively used in computational biology, for instance to understand diseases’ pathogenesis [23] and to show the implication of protein networks topology in genetics, personal genomics and therapy [35], among many other contexts.

By studying the topology of the human interactome and the ranking of nodes with respect to specific centrality measures, we aimed at unveiling if genes that are considered essential, *i.e.*, whose removal impact on a given organism’s ability to survive, can be associated to some specific structural feature. Differently from previous works, we focused the analysis on the pairwise connectivity of the network to understand if and how such essential genes (EGs) are related to this specific role in the network.

In this perspective, we first implemented a Critical Node Heuristic (CNH), based on the proposals made in [26] [41], that can be successfully applied to very large networks (billions of edges). Then we compared the performance of CNH against other four heuristics based on well know centrality measures, *i.e.*, Degree centrality (DC), Closeness centrality (CC), Betweenness Centrality (BC) and PageRank (PR). Finally, we performed a functional enrichment analysis to understand if there are common or distinct traits in the set of EGs found by the different centrality measures.

### A. Related works

A sizeable part of the research on PPI networks focused on finding genes of interest by means of methods and techniques borrowed from graph theory. With the aim of identifying the most essential genes, Jeong *et al.* [27] analysed the PPI network of *S. cerevisiae* showing that the degree distribution follows a power law. They suggested that hubs in the network are more likely to be essential with respect to lower degree nodes. This study has been followed by many others [21], [14], [29], [50] which tested the so called centrality-lethality hypothesis (the correlation between the centrality of a node in biological network and its essentiality) using several measures, mostly borrowed from the study of social networks [17]. Novel measures have been introduced with the aim of maximizing the number of essential genes discovered, by combining topological with biological information [46]. More recently Raman *et al.* [43] compared the percentage of essential genes found by different centrality measures in 20 PPI networks. They reported degree and betweenness as the best measures to find essential genes while closeness had poor performances.

There are also many studies that have criticized and/or revisited the centrality-lethality assumption. In [11] the authors observed that some previous works used biased datasets: on average, essential genes are more studied than non-essential, which can unbalance then number of known interactions between the two categories. In particular, they reported similar average degrees for both essential and non-essential genes in the PPI network of *S. cerevisiae*.

Brandes *et al.* [44] noticed that in previous works the poor performance of the closeness centrality was due to a misapplication of the measure to disconnected networks. They replicated the study and found that closeness has the better performance in identifying essential genes. Moreover, they pointed out how ranking by centrality strongly depends on the derivation of the network from the original data, *i.e.*, by the threshold used to determine if there is an interaction between two proteins. Our results on the closeness are not affected because we considered a single component.

Popatov *et al.* in [42] introduced the pairwise disconnectivity index as a new measure to quantify the importance of nodes/edges in regulatory networks. The differences with our approach are the following: a) they considered a pair of nodes *u, v* connected if there is at least one path between the nodes, *i.e.*, *u* → *v* or *v* → *u*, while we consider the pair connected only if there are both paths between the nodes, *i.e.*, *u* → *v* and *v* → *u*. The definitions are equivalent for undirected graphs; b) in [42] the connectivity of each node *v* is computed when node *v* is removed from the graph, in contrast we iteratively remove nodes with higher connectivity; c) they used the index to evaluate the centrality of nodes in regulatory networks, while we use it to study the connectivity of the graph; d) finally, their index was developed specifically to study regulatory networks having a limited number of nodes, while our heuristic can manage millions of vertices and billions of edges because it has a linear time complexity. In [43] they also tested the effectiveness of the pairwise disconnectivity as defined in [42] reporting that is not a good indicator for gene essentiality.

In [5], Boginski *et al.* introduced critical nodes as a technique to identify important nodes in biology networks. They developed a theoretical framework and some heuristics to solve the CNP. Unfortunately, these methods can be applied only on very small graphs. Moreover, they did not actually draw any topological and/or biological conclusion on the CNs found by the heuristics.

It is worth noting that other works, like [38] applied different methods to study the resilience of networks, mostly by measuring the change in the characteristic shortest path by removing nodes with high degrees. Finally, in [1] a graph of the human Interactome with similar global properties as the one in this work has been used to to discover disease pathways.

## II. Methods

### A. Centrality Measures

We used four centrality measures, *i.e.*, Degree centrality (DC), Closeness centrality (CC), PageRank (PR) and, Betweenness Centrality (BC). Each of them aims at capturing a different feature of the network’s nodes: a) DC selects the nodes with greatest degree in the network, *i.e.*, those that have the highest number of neighbours; b) CC ranks high the nodes that are the closest to all other nodes, *i.e.*, those nodes whose average distance to all other nodes of the network is minimal; c) PR, a recursive measure, selects the nodes pointed by other important nodes, *i.e.*, those having in turn high PR; d) finally, BC ranks high the nodes that are more often part of a shortest path of the network [16].

### B. Critical Nodes

In the CNP, the connectivity of a network is defined as the number of connected node pairs. More formally, let *G* = (*V, E*) be a graph, and let *C*_1_, *C*_2_, …, *C*_*N*_ be its connected components. The connectivity *f* (*G*) of *G*, is defined as:

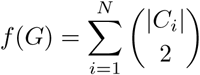

It is to be noted that *f* (*G*) equals the number of vertex pairs in *G* that are connected. The CNP is thus about finding a set of nodes *S* = {*n*_1_, *n*_2_, …, *n*_*k*_} of size *k*, namely the Critical Nodes of *G*, whose removal minimizes the connectivity of *G*. As already mentioned, the CNP is proven to be NP-complete [3], thus we needed to apply a heuristic approach for the analysis of the graph of interest.

The CN heuristic used here is based on the algorithm recently proposed by Paudel *et al.* [41] that optimally solves the CNP for *k* = 1, i.e. |*S*| = 1. We hereinafter refer to such heuristic as the Critical Node Hueuristic (CNH). The CNH is based on the framework proposed in [19] which uses the notions of Dominators [9], Strong Articulation Points [26] and Loop Nesting Forest [47] to build a novel data structure which stores the size of all Strongly Connected Component created by each node removal.

As showed in [41] the CNH outperforms other heuristics for CNs detection, resulting the best choice for solving the CNP in practice. Indeed, the authors experimentally evaluated their proposal on five different types of graph: Road networks, Peer to Peer networks, Web Graphs, Social networks, and Production Co-purchase networks. It is to be noted that none of them is related to the biological field.

In the present study, we used the mentioned five heuristics on the human interactome to identify the genes whose removal impact most on the interactome topology and connectivity, and mapped them to the essential gene sets in order to test the heuristics’ capability of identifying essential genes. In practice, we considered the same number of genes identified by each heuristics (*e.g.*, the first 1000, 2000, 4000 in the ranking for each heuristics) and calculated the intersection between them and the available EG datasets to understand the relation between the various measures and the essentiality of genes. Analysing the sets of genes selected by each heuristic, we were able to i) calculate how many CNs are EGs (or viceversa); ii) identify the best centrality measure to select EGs; iii) determine if CNs are related to EGs; iv) determine if EGs possess features that are not captured by CNs.

### C. Human protein-protein interaction network

We gathered human protein-protein interaction data from the BioGRID database, a biological interaction repository with data compiled through comprehensive curation efforts [40]. The network is composed of 17793 human proteins and 373123 experimentally validated interactions, obtained by the last available BioGRID *H. Sapiens* interaction dataset at the time of the analysis (Oct. 2019, v.3.5.177, namely BIOGRID-ORGANISM-Homo_sapiens-3.5.177.tab2.txt).

**TABLE I.**
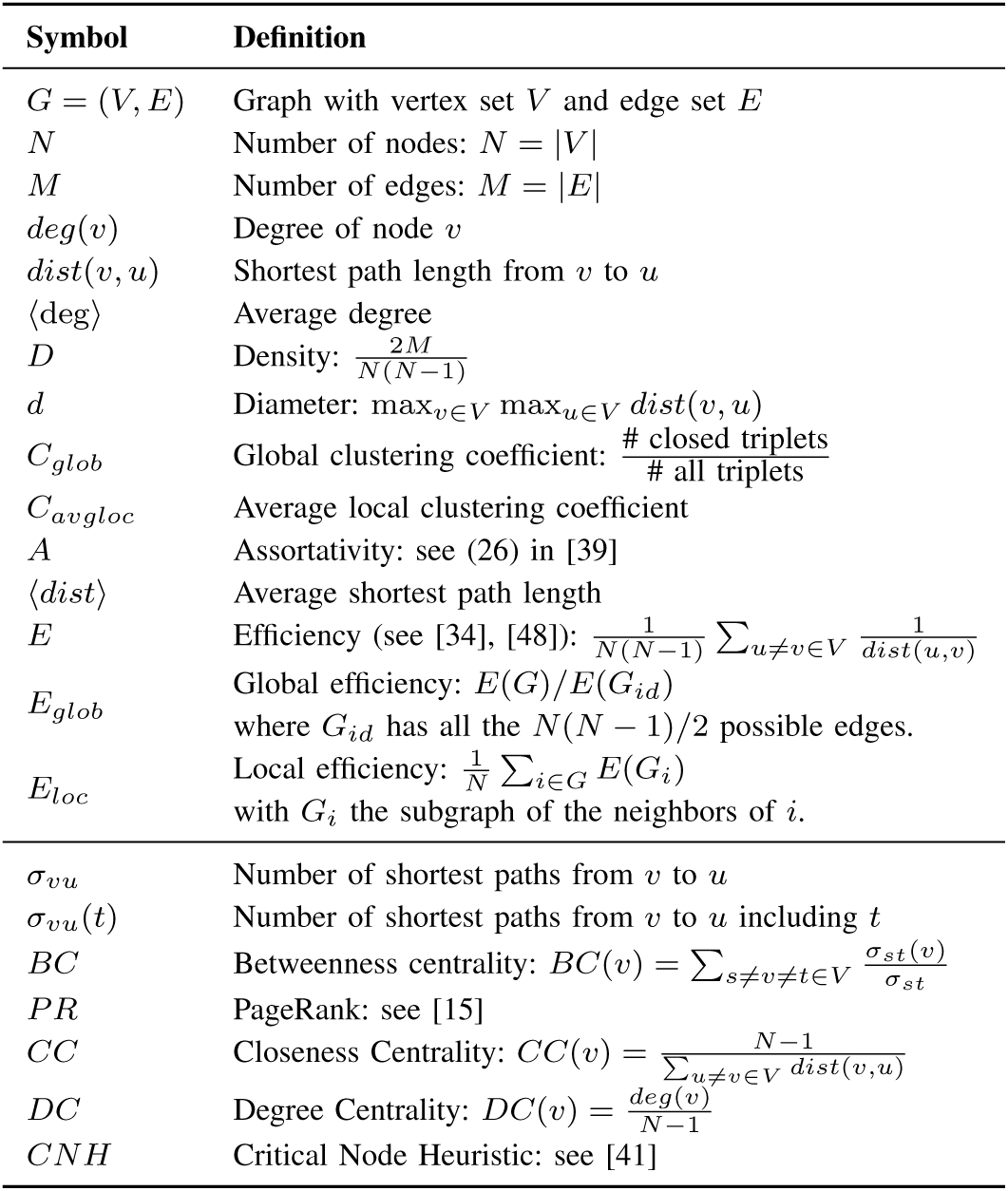
Notations and definitions used throughout the paper.

### D. Essential genes datasets

Simpler organisms (*e.g.*, single-cell) can often tolerate inactivating mutations in the majority of genes, while human cells generally require *more* genes essential for survival, likely because of increased complexity. Relevant efforts have been spent to identify the minimum set of human genes required for cell survival [6]. Luo *et al.* [18] used mutagenesis and high-throughput sequencing to identify prokaryotic and eukaryotic essential genes under diverse conditions and a revised essential-gene concept that includes all essential genomic elements. The authors collected such essential genes datasets in a freely available online resource, *i.e.*, the Database of Essential Genes (DEG). In *Homo sapiens*, 8256 genes are tagged as ‘essential’ in DEG (hereinafter “DEG” dataset). Similarly, Blomem *et al.* [4] used extensive mutagenesis in two haploid human cell types (HPA1 and KBM7 cell lines) to identify approximately 2000 genes which seem to be required for viability and optimal fitness under *in vitro* conditions. In details, 2180 genes are identified as essential in the HAP1 cell line, and 2053 in the KBM7 cell line. The essential genes in common between these two cell lines are 1734 (intersection), while the union (hereinafter “HK” dataset) counts 2501 genes.

### E. Gene functional enrichment analysis

For space reasons, only three of the five heuristics have been selected for comparison, namely CNH, CC and DC. The CNH is the best in finding CNs, whereas CC and DC are the best in selecting EGs for both datasets, as discussed in Subsection III-B. The functional enrichment analysis on genes found by the three algorithms on two independent datasets (DEG and HK; it is to be noted that HK is almost entirely a subset of DEG) has been performed with the functional enrichment platform Enrichr [10], [32]. GO biological processes with an adjusted p-value *<* 0.05 have been considered for downstream analyses. Enriched terms found with Enrichr have then been processed using the REVIGO tool [45], which allowed us to summarize gene ontologies by removing redundant ones. Analysis with REVIGO has been performed with the default suggested parameters (*i.e.*, medium allowed similarity between terms (0.7), adjusted p-value derived from Enrichr analysis associated to each GO term, whole UniProt as reference database with GO term size and SimRel as semantic similarity measure to be used).

## III. Discussion

### A. Graph Structure

The graph is composed by six connected Component and fifteen isolated nodes. Five components include just 2 nodes, while the sixth is a large component that comprises the 0.99% of vertices. Hereafter, all statements to and analyses of the Interactome are meant to be referred to its largest component.

Table II summarizes the values of the global measures of the graph. It is important to notice that the average (or characteristic) path length 〈*dist*〉 is quite small, *i.e.*, ~3, and the interactome diameter (the longest of the shortest paths) is equal to 8. This value is in accordance to the formula *ln*(*N*)/*ln*(*k*) which holds for Erdos-Renyi (ER) random graphs. The value of *C*_*glob*_, on the other hand is nearly 30 times that of an ER graph with the same nodes/edges (*C*_*ER*_ = 0.0022). These results permit to classify the network as Small-World.

**TABLE II.** Global measures of the graph

| Measure | Value | Measure | Value | Measure | Value |
| --- | --- | --- | --- | --- | --- |
| N | 17793 | $\langle \text{deg} \rangle$ | 38.621 | A | -0.0941 |
| M | 343115 | $\langle \text{dist} \rangle$ | 2.9667 | $E_{glob}$ | 0.35322 |
| d | 8 | $C_{avgloc}$ | 0.1213 | $E_{loc}$ | 0.50238 |
| D | 0.0022 | $C_{glob}$ | 0.0635 | | |

We also note that the value of the Assortativity (−0.09) implies that there is no correlation between nodes degree, a node with high degree has the same probability to bond with a node with a similar degree or with a lower degree for different biological networks. Given the degree distribution, most of the high degree nodes must have more interactions with lower degree nodes. This result is in agreement with those reported by [43], and [39].

We analyzed the degree distribution and fit the data against a power law and a log-normal distribution. The analysis has been carried out by mean of the `power_law` python package [2]. By testing both distributions, varying the critical degree (*k*_*c*_) from which the computation of the fit was carried out, and comparing the results using the normalized likelihood ratio we found that in all cases the log-normal distribution should be preferred. Figure 1 reports the result for *k*_*c*_ = 5 (which includes 75% of the nodes). In a recent paper, Broido *et al.* [8], by analysing 1000 networks, empirically demonstrated that the log-normal distribution is more frequently the best fit (more than 3 times favoured with respect to the power law). Many works described the power law distribution as a general feature of biological networks [20], [24], [27], [49] while others rise a number of reservations [30], [36]. While power law implies the existence of hubs and the so called scale-free topology, the log-normal distribution has a well-defined mean and variance. Our observation that the current state human Interactome’s topology fits with a log-normal distribution, if confirmed, suggests the existence of significant investigative biases or that available data can still be incomplete, as already observed [37], among other issues.

**Fig. 1.**
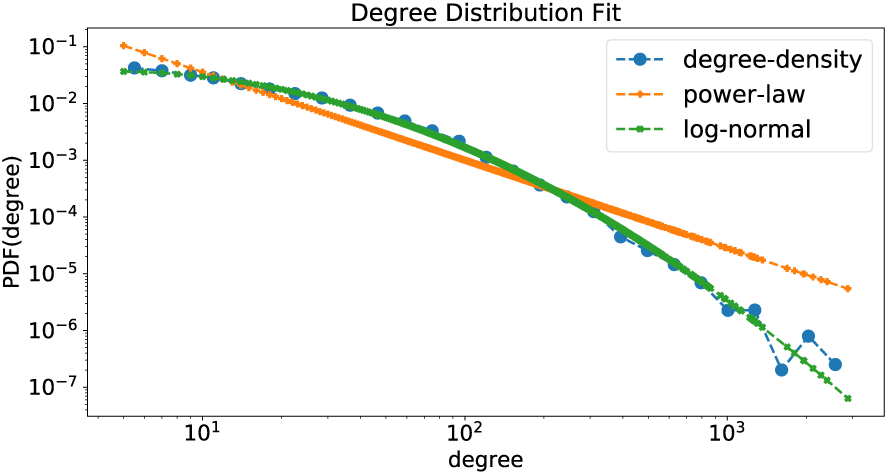
The degree distribution is shown along with the fit obtained by a power law and a log-normal functions. The fit starts at *k*_*c*_ = 5. It is clear that the log-normal distribution better approximate the underlying data.

As we have analyzed just one network we cannot draw any general conclusion: the degree distribution strongly depends on the topology (actually is what defines the topology) which, in turns, strongly depends on how the network is reconstructed and/or sampled. Most of the complex networks cannot be directly measured, but rather they are built on some knowledge or sampled from larger data. Each technique introduces its own bias which is hard to rule out from the analysis. We have intentionally reported a large number of topological features to make it easier to compare future analysis on different (hopefully more accurate) PPI networks with our results.

### B. Network Connectivity

In this subsection we discuss how CNs removal affect graph’s connectivity depending on the applied heuristic and the relation between CNs and EGs. As shown in Figure 2 (and as expected), the CNH is the best heuristic for CNs detection: it reduces the connectivity of the graph faster than the other four heuristics we used. Furthermore, as show in Table III, CNH is almost 2× faster than other heuristics even considering that we computed CNH rank after each node removal, while for the other heuristics (BC, CL, DG, and PG) we computed the node ranking only once, via the *igraph* C library [12].

**TABLE III.**
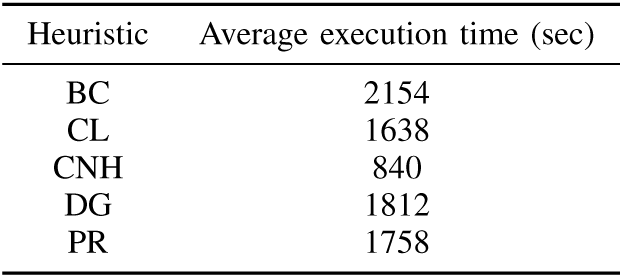
Average execution time of the five heuristics on centOS machine with 2 intel(R) xeon(R) CPU E5-2683 v4 2.10GHz and 512GB of memory

**Fig. 2.**
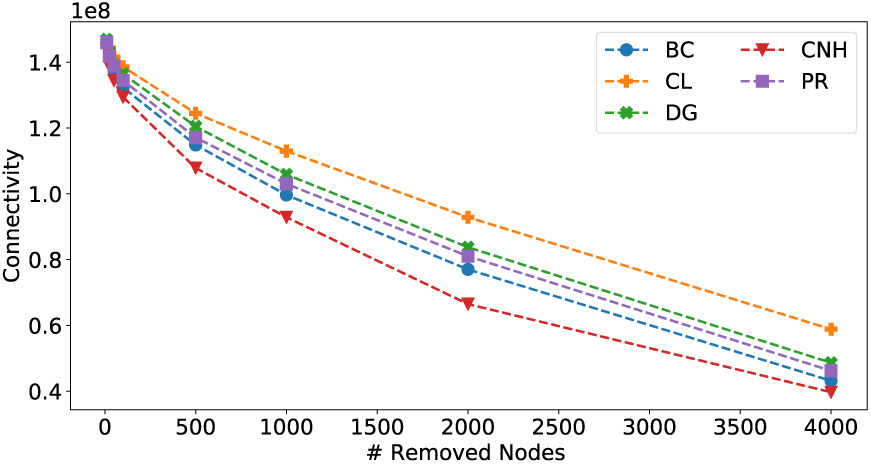
Graph connectivity trend respect to nodes removal

From our results we observed that CC and DC are the worst heuristics for CNs detection, as expected. Indeed, CC and DC are better suited to identify nodes used to diffuse information in the network instead of reducing network connectivity [7].

Given the performance of the five heuristics in identifying CNs, we analysed the sets of genes selected by each heuristic. In particular, we checked how many of such genes are EGs. As baseline for our analysis, we compared the sets selected by each heuristic with the sets selected by a Random procedure and an Optimal procedure. The percentage of EGs found by the Random procedure varies depending on the dataset considered (DEG or HK). Specifically, the number of EGs selected by a Random procedure can be computed using the mean of the Hypergeometric distribution as the number of extractions varies. On the other hand the sets of genes selected by the Optimal procedure contain only EGs. The results of our analysis are summarized in Tables IV, V and VI.

The percentage of EGs found by each heuristic was always greater than the baseline set by the Random procedure, in both datasets (see Figures 3, 4, 5, 6). None of the heuristics is able to select only EGs, but some of them perform better than the others. In particular, CC results to be the best centrality measure for selecting EGs. In both dataset the percentage of EGs selected by CC is higher than all other heuristics for any set size. On the other hand CNH is the worst choice if we are looking for EGs. Moreover, for the HK dataset the CNH is only slightly better than the Random procedure, especially as the size of selected genes increases.

**Fig. 3.**
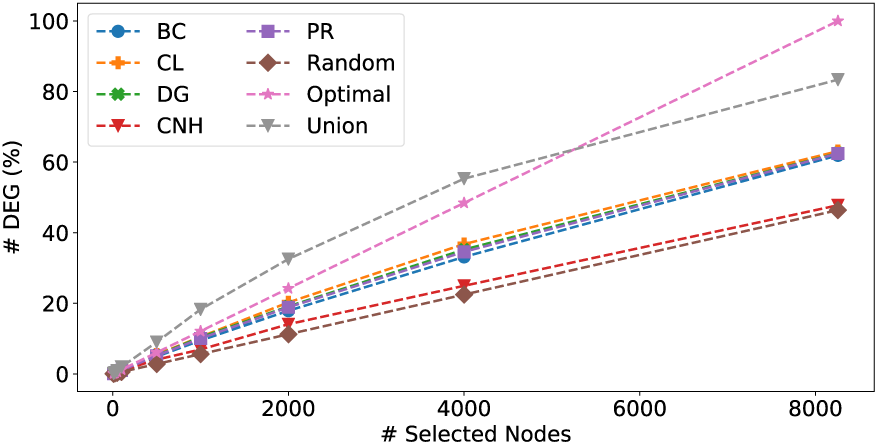
[DEG dataset] Percentage of selected EGs.

**Fig. 4.**
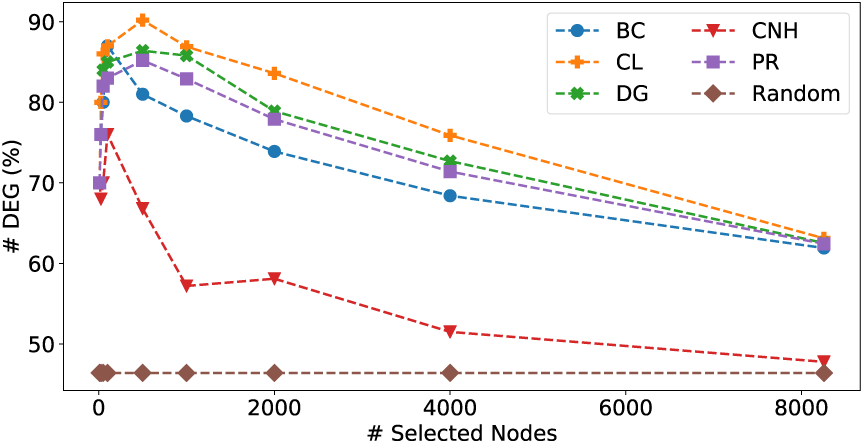
[DEG dataset] Percentage of EGs inside each set.

**Fig. 5.**
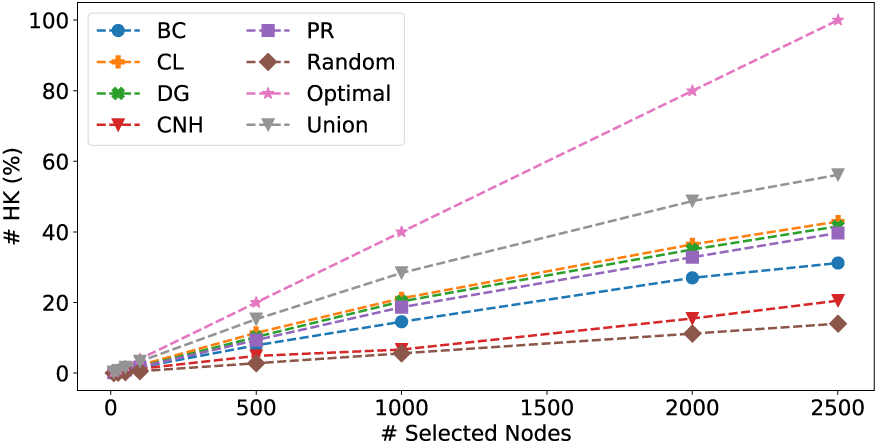
[HK dataset] Percentage of selected EGs.

**Fig. 6.**
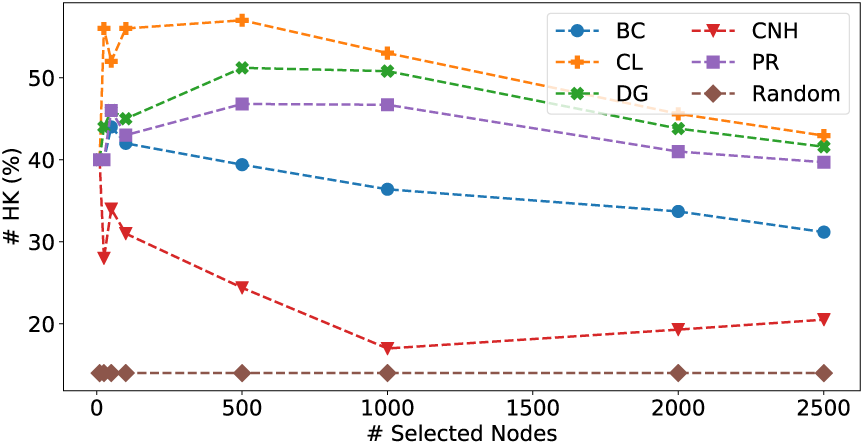
[HK dataset] Percentage of EGs inside each set.

Our results clearly show that only a part of EGs are central nodes in the PPI network we have analyzed. Closeness centrality and degree centrality are distinct features of nodes that have many contacts and can quickly reach many other nodes. It is worth noting that closeness centrality has a low power of discriminating among nodes because, by definition, the permitted values are packed in a small interval.

The most interesting result, related to the connectivity, is that the heuristics that better disrupt the network are those that find less EGs, for the same set of genes selected. This behaviour is clearly shown in Figures 2, 3, 4, 5, 6.

We conclude that EGs are not fundamental nodes for the network connectivity, thus, only a small portion of EGs (especially for the HK dataset) are responsible for holding together different parts of the networks, and increasing the number of connected genes pairs. In the next section we try to give a biological interpretation for these EGs that are also CNs.

It is worth noting that in figure 4, for all the heuristics, the percentage of discovered EG decays as the number of selected genes increases, a characteristic behaviour which has been observed also in [43] for different kind of PPI networks. We analysed also the union of the genes selected by each heuristic and we observed that neither with the union we are able to select all the EGs. This observations lead us to the conclusion that centrality measures cannot solve the problem of identifying EGs, only a portion of them can be selected by the heuristics.

**TABLE IV.** [DEG dataset] number of EGs selected, percentage of EGs selected respect to the set size and percentage of EGs selected respect to the total number of EGs in the network by each heuristic.

| #Nodes | CNH | BC | PR | CL | DG |
| --- | --- | --- | --- | --- | --- |
| 10 | 7 (70% \| 0.08%) | 7 (70% \| 0.08%) | 7 (70% \| 0.08%) | 7 (70% \| 0.08%) | 7 (70% \| 0.08%) |
| 25 | 17 (68% \| 0.20%) | 19 (76% \| 0.23%) | 19 (76% \| 0.23%) | 20 (80% \| 0.24%) | 19 (76% \| 0.23%) |
| 50 | 35 (70% \| 0.42%) | 40 (80% \| 0.48%) | 41 (82% \| 0.49%) | 43 (86% \| 0.52%) | 42 (84% \| 0.50%) |
| 100 | 76 (76% \| 0.92%) | 87 (87% \| 1.05%) | 83 (83% \| 1.00%) | 87 (87% \| 1.05%) | 85 (85% \| 1.02%) |
| 500 | 334 (66.8% \| 4.04%) | 405 (81% \| 4.90%) | 426 (85.2% \| 5.15%) | 451 (90.2% \| 5.46%) | 432 (86.4% \| 5.23%) |
| 1000 | 572 (57.2% \| 6.92%) | 783 (78.3% \| 9.48%) | 829 (82.9% \| 10.04%) | 869 (86.9% \| 10.52%) | 858 (85.8% \| 10.39%) |
| 2000 | 1163 (58.1% \| 14.08%) | 1478 (73.9% \| 17.90%) | 1559 (77.9% \| 18.88%) | 1673 (83.6% \| 20.26%) | 1579 (78.9% \| 19.12%) |
| 4000 | 2061 (51.5% \| 24.96%) | 2737 (68.4% \| 33.15%) | 2856 (71.4% \| 34.59%) | 3037 (75.9% \| 36.78%) | 2909 (72.7% \| 35.23%) |
| 8256 | 3945 (47.78% \| 47.78%) | 5112 (61.91% \| 61.91%) | 5155 (62.43% \| 62.43%) | 5211 (63.11% \| 63.11%) | 5163 (62.53% \| 62.53%) |

**TABLE V.** [HK dataset] Number of EGs selected, percentage of EGs selected respect to the set size and percentage of EGs selected respect to the total number of EGs in the network by each heuristic.

| #Nodes | CNH | BC | PR | CL | DG |
| --- | --- | --- | --- | --- | --- |
| 10 | 4 (40% \| 0.15%) | 4 (40% \| 0.15%) | 4 (40% \| 0.15%) | 4 (40% \| 0.15%) | 4 (40% \| 0.15%) |
| 25 | 7 (28% \| 0.27%) | 10 (40% \| 0.39%) | 10 (40% \| 0.39%) | 14 (56% \| 0.55%) | 11 (44% \| 0.43%) |
| 50 | 17 (34% \| 0.67%) | 22 (44% \| 0.87%) | 23 (46% \| 0.91%) | 26 (52% \| 1.03%) | 23 (46% \| 0.91%) |
| 100 | 31 (31% \| 1.23%) | 42 (42% \| 1.67%) | 43 (43% \| 1.71%) | 56 (56% \| 2.23%) | 45 (45% \| 1.79%) |
| 500 | 122 (24.4% \| 4.87%) | 197 (39.4% \| 7.87%) | 234 (46.8% \| 9.35%) | 285 (57% \| 11.39%) | 256 (51.2% \| 10.23%) |
| 1000 | 167 (16.7% \| 6.67%) | 364 (36.4% \| 14.55%) | 467 (46.7% \| 18.67%) | 530 (53% \| 21.19%) | 508 (50.8% \| 20.31%) |
| 2000 | 387 (19.3% \| 15.47%) | 675 (33.7% \| 26.98%) | 821 (41% \| 32.82%) | 912 (45.6% \| 36.46%) | 876 (43.8% \| 35.02%) |
| 2501 | 513 (20.51% \| 20.51%) | 780 (31.18% \| 31.18%) | 993 (39.7% \| 39.7%) | 1074 (42.94% \| 42.94%) | 1040 (41.58% \| 41.58%) |

**TABLE VI.** [DEG and HK datasets] Number of EGs selected, percentage of EGs selected respect to the total number of EGs in the network and standard deviation of the Random procedure.

| #Nodes | R ($\mu$) | R ($\sigma$) | #Nodes | R ($\mu$) | R ($\sigma$) |
| --- | --- | --- | --- | --- | --- |
| 10 | 4.64 (0.05%) | 1.58 | 10 | 1.4 (0.05%) | 1.1 |
| 25 | 11.6 (0.14%) | 2.49 | 25 | 3.5 (0.14%) | 1.73 |
| 50 | 23.2 (0.28%) | 3.52 | 50 | 7 (0.28%) | 2.45 |
| 100 | 46.4 (0.56%) | 4.97 | 100 | 14 (0.56%) | 3.47 |
| 500 | 232 (2.81%) | 10.99 | 500 | 70 (2.81%) | 7.66 |
| 1000 | 464 (5.62%) | 15.32 | 1000 | 140 (5.62%) | 10.68 |
| 2000 | 928 (11.24%) | 21.01 | 2000 | 280 (11.24%) | 14.64 |
| 4000 | 1856 (22.48%) | 27.77 | 2501 | 351 (14%) | 16.11 |
| 8256 | 3831 (46.4%) | 33.17 | - | - | - |

### C. Biological interpretation

The functional enrichment analysis performed on the DEG dataset revealed a different set of terms found with the three distinct algorithms (DC, CC, CNH). Globally, we found a good intersection between the three algorithms (41.9% of overlapping terms, Figure 7). While the genes found using CC and DC showed a relevant intersection of GO terms (19.4% between them) and an identical summarized REVIGO output (Figure 8) among themselvels, the CNH found a relevant set of genes (17.1%) which is unique. As a consequence, the summary of the GO terms of CHN revealed a set of different gene ontologies in comparison to CC and DC but that are very close to those of CC and DC (regulation of transcription, localization of proteins to the membrane and viral dynamics), but, interestingly, also a set of processes like autophagy, ventricular septum development and mitotic phase of the cell cycle which are unique to the CHN EG dataset. The same analysis on GO cell component terms highlighted that the three algorithms led to an enrichment in ontologies related to the generic cytoplasm compartment, while showing their own peculiarities (Figure 9). While CC and DC had some overlapping terms like focal adhesion and axon, CNH showed a completely different set of terms, such as “aggresome” and “membrane raft”. Moreover, other found terms look more detailed in comparison to the others (*e.g.*, “dendrite” and “neurotransmitter receptor complex” in comparison to “axon”, or “perinuclear region of cytoplasm” in comparison to a generic “cytosolic part”).

**Fig. 7.**
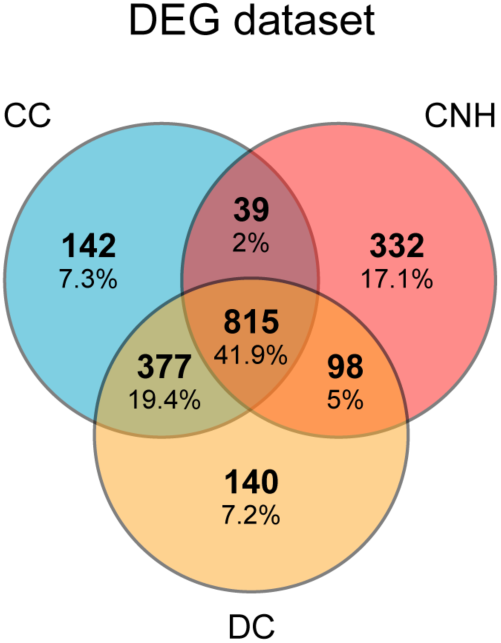
Functional enrichment analysis of the DEG essential gene sets found with the utilized heuristics. Venn diagram showing the intersection between enriched GO biological processes for CC (blue), DC (yellow), CNH (pink) heuristics.

**Fig. 8.**
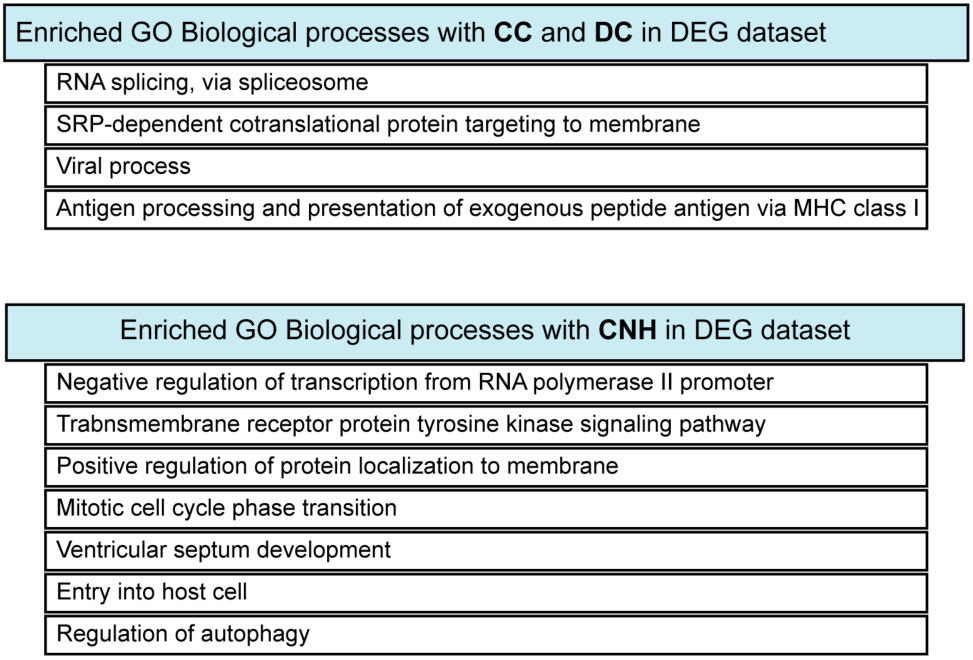
Functional enrichment analysis of the DEG essential gene sets found with the utilized heuristics. List of top enriched GO biological processes (adjusted p-value<0.05) for CC and DC (top table) and CNH (bottom table).

**Fig. 9.**
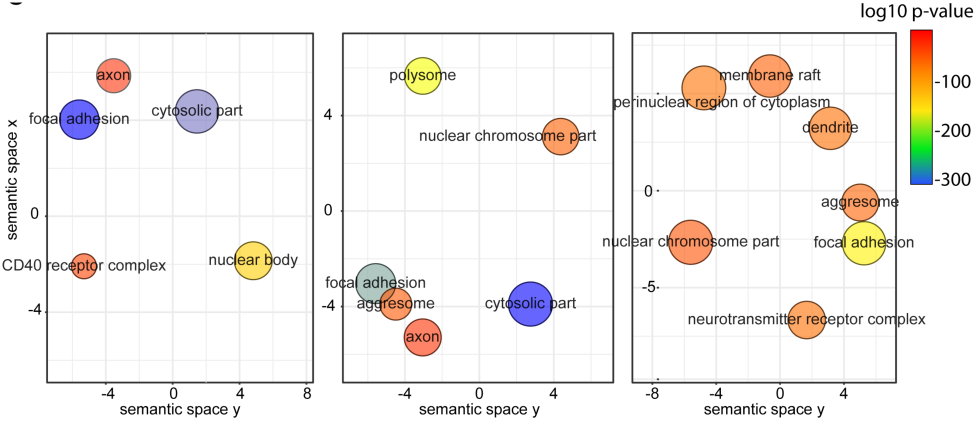
Functional enrichment analysis of the DEG essential gene sets found with the utilized heuristics. Scatterplot for GO cellular component terms derived from REVIGO analysis for CC (left panel), DC (central panel) and CNH (right panel). Node size is mapped on the log size parameter; node color is mapped on log10 p-value.

## IV. CONCLUSION

We analyzed with a graph theoretical approach the features of the human interactome focusing on the connectivity of the network. Our results show that, while EGs are more easily found by a genes ranking based on CC and DC, those are not the nodes whose removal impact the most the connectivity of the network. In other words, EGs removal decreases significantly the organism’s survival, but it has a minor impact on disrupting the PPI connectivity respect to the nodes identified by the CNH. Moreover, centrality measures seems to fail in identifying all EGs which implies that some of them are not central in the network and different techniques should be applied to discover them. Considering the genes found with CNH, we performed a functional enrichment analysis, which highlighted an enrichment of gene ontology terms that are not found when performed with the DC or CC-related gene sets. This will prompt us to make further investigations, linking the computational theory behind CHN to a valuable biological meaning.

Our analysis is meant to be a first step in understanding the feature of PPI networks. As a future work we plan to study how the average shortest path length vary under the removal of CNs and EGs. Moreover, we want to test our methods on a large set of biological networks to understand if our findings can be generalized. Another appealing direction that should be explored regards the sub-network induced by essential genes: some works [14] reported high level of clusterization among them, but there are no study about the connectivity of such network.

## Footnotes

1 We note that this could be also referred as the Critical Node Detection Problem (CNDP), of which CNP is a specific instance where the metric for measuring the connectivity is the pairwise connectivity.

